# Growth of brown trout in the wild predicted by embryo stress reaction in the laboratory

**DOI:** 10.1101/2022.07.07.499115

**Authors:** Jonas Bylemans, Lucas Marques da Cunha, Laetitia G. E. Wilkins, David Nusbaumer, Anshu Uppal, Claus Wedekind

**Author notes:** Co-First authors (Jonas Bylemans, Lucas Marques da Cunha).

## Abstract

Laboratory studies on embryos of salmonids, such as the brown trout (*Salmo trutta*), have been extensively used to study environmental stress and how responses vary within and between natural populations. These studies are based on the implicit assumption that early life-history traits are relevant for stress tolerance in the wild. Here we test this assumption by combining two datasets from studies on the same 60 full-sib families. These families had been experimentally produced from wild breeders to determine, in separate samples, (i) stress tolerances of singly kept embryos in the laboratory and (ii) growth of juveniles during 6 months in the wild. We found that growth in the wild was well predicted by larval size of their full sibs in the laboratory, especially if these siblings had been experimentally exposed to a pathogen. Exposure to the pathogen had not caused elevated mortality among the embryos but induced early hatching. The strength of this stress-induced change of life history was a significant predictor of juvenile growth in the wild: the stronger the response in the laboratory, the slower the growth in the wild. We conclude that embryo performance in controlled environments can be useful predictors of juvenile performance in the wild.

## Introduction

Salmonids are not only charismatic fish of high ecological and socio-economical relevance but also excellent models for experimental research, especially on early developmental stages. External fertilisation and the lack of parental care allow for full-factorial *in vitro* fertilizations under controlled conditions. Embryos can then be raised in separated groups or even singly in multi-well plates, for example, to estimate the genetic and the maternal environmental effects on embryo mortality (Houde et al. 2013, Houde et al. 2016), to study developmental problems (Evans and Neff 2009), or to characterise symbiotic microbial communities (Wilkins et al. 2016). Embryo performance can then be linked to parental characteristics to learn more about, for example, the information content of sexual signals (Neff and Pitcher 2005, Wedekind et al. 2008, Huuskonen et al. 2011, Janhunen et al. 2011), the genetic quality of dominant males (Jacob et al. 2007), or fitness consequences of different female reproductive strategies (Jacob et al. 2010, Kekäläinen et al. 2010). Embryo performance can also be studied under different environmental conditions. Large numbers of independent replicates then allow estimating the relevance of environmental factors and the evolutionary potential of populations to adapt to them. The latter is typically revealed in 2- or 3-way interactions between parental and environmental factors on embryo performance. Environmental factors that have been studied on embryos of experimentally produced families include increased temperatures in the context of climate change (Burt et al. 2012, Muñoz et al. 2014), different types of chemical pollutants (Marques da Cunha et al., 2019; Nusbaumer et al., 2021b), pollution by nanoparticles (Clark et al., 2016; Yaripour et al., 2021), organic pollution (Wedekind et al. 2010, Nusbaumer et al. 2021a) and pathogens (von Siebenthal et al. 2009, Pompini et al. 2013, Clark et al. 2014, Wilkins et al. 2017). Full-factorial crosses can even be used to estimate the variance components of commercial traits in farmed salmon (Colihueque, 2010; Ødegård et al., 2011) or to study effects of sperm cryopreservation (Nusbaumer et al. 2019). Factorial breeding within and between populations and monitoring of embryo performance has also been used to study the causes of phenotypic differentiation among populations (Aykanat et al. 2012) or to determine hybrid vigour as indicator of inbreeding depression (Clark et al. 2013a, Stelkens et al. 2014).

Most studies that are based on factorial breeding designs have focused on embryos or larvae and have ignored later developmental stages, with few exceptions: Juveniles that resulted from experimental breeding have been raised in captivity to study genotype-by-environmental effects (Evans et al. 2010), genetic aspects of life history (Forest et al. 2016), maternal environmental in different temperature environments (Thorn and Morbey 2018), or effects of enriched vs non-enriched environments (Yaripour et al. 2020). Studies on juveniles released into the wild and recaptured later are scarce and, so far, generally suffer from low sample size because of low recapture rates (Wedekind et al. 2008, von Siebenthal et al. 2017). Consequently, the conclusions drawn from laboratory studies on experimentally produced families assume that the observed reactions reveal effects that are also relevant in the wild. This assumption is still poorly backed up.

Here we combine data of two studies that could be successfully performed in separate samples from the same 60 full-sib families. These families had been experimentally produced in two full-factorial breeding blocks using gametes collected from wild-caught males and females. The breeders had been sampled from a stream that represents a mostly pristine environment within the Swiss Plateau (Marques da Cunha et al. 2019, Nusbaumer et al. 2021b) and a population that shows no signs of elevated inbreeding (Clark et al. 2013a, Stelkens et al. 2014). One sample of freshly fertilized eggs per each of the 60 families was used in a controlled laboratory environment to study embryo growth and stress tolerance (Wilkins et al. 2017). The other sample was incubated under hatchery conditions and stocked into a natural streamlet at early larval stages following routine procedures of the local fishery authorities. Approximately six months later, a significant number of these fish could be successfully sampled at the end of their first summer, as revealed by molecular markers. Parental assignments and molecular sexing could then be applied to study inbreeding depression in the wild (Bylemans et al. 2023). Here we combine the findings on the embryos (Wilkins et al. 2017, 2018) with the findings on the juveniles (Bylemans et al. 2023) to test how the outcome of laboratory studies on embryos correlate with juvenile performance in the wild.

## Material and methods

The experiments started with male and female brown trout being caught shortly before the spawning season from the *Rotache* stream (a tributary of the *Aare* river, see Stelkens et al. (2012) for a description of the genotypes and phenotypes of this and neighbouring populations). They were kept in the *Fischereistützpunkt Reutigen* and regularly checked for ovulation until the eggs of 12 females could be stripped and fertilized with milt of 10 males to create two full-factorial breeding blocks (6 × 5 each) on the same day. See Wilkins et al. (2017) for a detailed description of the procedure. Fin clips were collected and stored in 70 % ethanol at -20 °C for molecular analyses.

Prior to fertilization, total egg weight per female was determined and four eggs per female were frozen in liquid nitrogen for later analyses of the carotenoids astaxanthin, capsanthin, lutein, and zeaxanthin as described in Wilkins et al. (2017). After fertilisation and egg hardening (*>*2 hours), photos of each full-sib family (eggs in monolayer in individual Petri dishes) were taken to later count the eggs and determine the average egg weight per female (total egg weight / egg count).

A subset of 24 fertilized eggs per full-sib family was transported to the laboratory and raised singly (each egg in its own 2mL well of a 24-well plate) in a climate chamber under controlled experimental conditions to assess the effects of egg carotenoids on the tolerance of embryos to the bacterium *Pseudomonas fluorescens*. A bacterial strain was used that had previously been found to reduce embryo growth and affect hatching time but would not significantly elevate mortality (Clark et al. 2013a, Clark et al. 2013b). The details of the bacterial culture and exposure are provided in Wilkins et al. (2017). Briefly, half of the embryos per family received 0.2 mL standardized water (OECD 1992) with 2 × 10^6^ bacterial cells per each well to a final water volume of 2 mL/well, the other half received 0.2 mL standardized water only (sham controls). The treatment happened either 18 dpf (days past fertilization; 1^st^ breeding block) or 49 dpf (2^nd^ breeding block).

Shortly before hatching was expected to start, 8 embryos per family were sacrificed for a study on carotenoid consumption (Marques da Cunha et al. 2018). All remaining eggs were checked daily to record individual hatching time and to take photos of the freshly hatched larvae. These photos were used to determine hatchling length (L_hatching_) and hatchling yolk sac volume (YS_hatching_). Because hatchling size is likely to vary with the timing of hatching, larval length and yolk sac volume were again measured for each larva 14 dph (days past hatching; L_14dph_ and YS_14dph_, respectively). This could be done in a sample of 815 larvae (after 5 larvae whose measurements of larval growth per loss of yolk sac over these 14 days exceeded 3 standard deviations from the global mean had been excluded as outliers). From these measurements, larval length at 75 dpf, the day when all larvae had hatched (and 11 days after first hatching), was calculated as

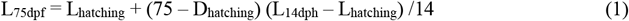

where D_hatching_ is the number of days from fertilization to hatching, and L_14dph_ the larval length at 14 dph. Yolk sac volume at 75 dpf was determined as

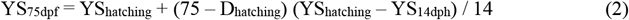

The remaining 1,925 eggs (mean±SD number per full-sib family = 32.1±14.8) were incubated from the day of fertilization under routine hatchery conditions in the cantonal *Fischereistützpunkt Reutigen* at constant 8.5°C and stocked into the *Mühlibach* streamlet (tributary to the *Rotache*; 46.804459°N, 7.690544°E) at 105 dpf, i.e., in early March at a late yolk-sac stage when emergence from gravel would usually happen and larvae would start exogenous feeding.

In late August (281 dpf, i.e., nearly 6 months after release into the wild), electrofishing was used to sample brown trout juveniles from the *Mühlibach* streamlet as reported in Bylemans et al. (2023). The fish were narcoticized (0.075 g/L tricaine methanoesulfonate buffered with 0.15 g/L NaHCO3) and photographed on a weighting scale to later extract fork length and weight. Fin clips were collected and stored in 70 % ethanol at 4 °C for molecular analyses.

Fin clips of adult breeders and a random subset wild-caught juveniles (375 of 518 juveniles) were used for microsatellite genotyping and genetic sexing as described in Palejowski et al. (2022). Briefly, the fish were genotyped at 13 microsatellite loci and a sex determination locus. Parental assignment was based on the full-likelihood approach implemented in Colony v2.0.6.5 (Jones and Wang, 2010) with a threshold of 0.98.

Statistical analyses were done in JMP Pro17 and R 4.0.2 (R Development Core Team, 2010). Standard F-tests were used to compare means. Linear mixed-effect models (LMM) were used to evaluate the relationship between laboratory-based measures (means per sib group and treatment) and the sizes of their full-sibs caught from the wild. Dam and sire identities were included as random effects. Treatment effects were tested using the mean difference in response variables (control – exposed to *P. fluorescens*) as predictors of juvenile size in the wild. Breeding block was added to the model when analysing correlations to hatching date, because the timing of the experimental exposure to *P. fluorescens* could potentially affect the timing of hatching. Collinearity between predictor variables was evaluated using Pearson correlation coefficients to avoid inclusion of highly correlated predictor variables (i.e. |r| > 0.5) within the same model (Dormann et al., 2013). Spearman correlation coefficients (r_s_) were used to test for correlations between recapture rates in the wild and embryo performance in the laboratory, based on means per parent (after averaging over sib-group families).

## Results

Juvenile body length was on average 7.1 times larger than larval body length at 75 dpf and was well predicted by mean larval length and mean yolk sac volume of their full sibs at 75 dpf (Table 1; Fig. 1a; Fig. S1). Juvenile body length was also positively correlated to mean egg weight (LMM; F = 8.9, p = 0.04; Table S1a; Fig. S2) but not significantly to the carotenoid content of the eggs (Table S1b).

**Table 1.** Linear mixed model on juvenile length as predicted by (a) larval length at 75 dpf (days past fertilization, i.e., about 6 months before the juvenile were caught), and (b) yolk sac volume per family at 75 dpf (using means per treatment and family each). Parental identities were included as random factors. Significant p-values are highlighted in bold.

| Effects | d.f. | F | Variance component | p |
| --- | --- | --- | --- | --- |
| *a) Larval length* |  |  |  |  |
| Fixed effects: |  |  |  |  |
| Larval length | 1 | 6.3 |  | **0.02** |
| Random effects<sup>1</sup>: |  |  |  |  |
| Dam |  |  | 20.0 ± 10.2 | **0.05** |
| Sire |  |  | 2.9 ± 3.1 | 0.35 |
| Residual |  |  | 81.1 ± 6.9 |  |
| *b) Yolk sac* |  |  |  |  |
| Fixed effects: |  |  |  |  |
| Yolk sac volume | 1 | 13.5 |  | **0.006** |
| Random effects<sup>1</sup>: |  |  |  |  |
| Dam |  |  | 13.0 ± 8.4 | 0.12 |
| Sire |  |  | 4.0 ± 3.5 | 0.26 |
| Residual |  |  | 81.3 ± 7.0 |  |

<sup>1</sup>REML unbounded variance components ± standard error, Wald p-values

**Figure 1.**
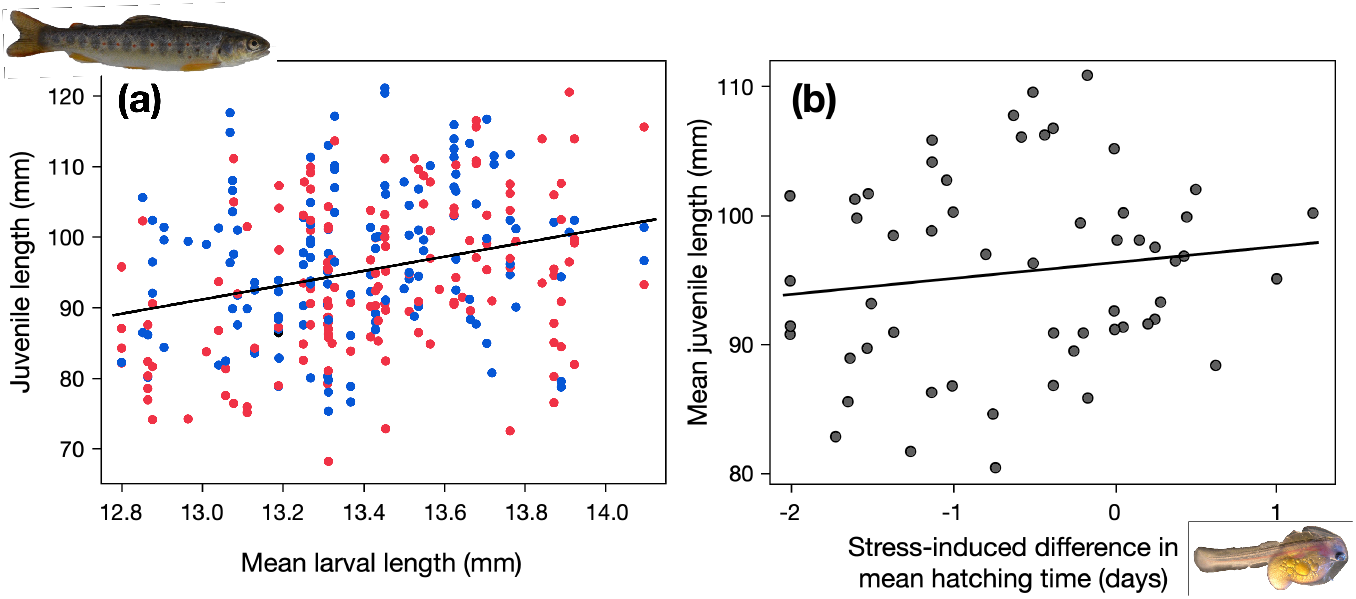
Juvenile length after 6 months in the wild predicted by larval characteristics in the laboratory. (a) Juvenile length *vs*. the mean larval length per full-sib family at 75 dpf (past last fertilization), for male (blue symbols) and female juveniles (red symbols). (b) Mean juvenile length per full-sib family *vs*. the stress-induced difference in mean hatching date of their full sibs in the laboratory. The regression lines are drawn to illustrate the direction of the correlations. See Table 1 for statistics that takes parental effects into account. The photos show a freshly hatched larvae still partly in its egg membrane (© Manuel Pompini) and an average-sized juvenile after its first spring and summer in the wild (© C. Wedekind).

When separately testing larvae that had or had not been stressed with *P. fluorescens*, the average days of hatching in the laboratory did not predict juvenile body length in the wild (Table S2). However, the experimental stress induced a change in mean hatching date that was a good predictor of growth in the wild: the stronger the stress response in the laboratory, the smaller the juveniles grew in the wild (Fig. 1b; Table 2). Adding breeding blocks and mean larval length as factors to the LMM did not change this conclusion (Table S3).

**Table 2.** Linear mixed model on juvenile length as predicted by the stress-induced difference in hatching date under laboratory conditions, i.e., the difference between the mean hatching date under control conditions and the mean hatching date after exposure to *P. fluorescens*. Parental identities were included as random factors in all models. Significant p-values are highlighted in bold.

| Effects | d.f. | F | Variance component | p |
| --- | --- | --- | --- | --- |
| Fixed effects: |  |  |  |  |
| Difference in hatching date | 1 | 5.4 |  | **0.02** |
| Random effects<sup>1</sup>: |  |  |  |  |
| Dam |  |  | 30.4 ± 14.5 | **0.04** |
| Sire |  |  | 3.4 ± 2.9 | 0.23 |
| Residual |  |  | 79.0 ± 6.7 |  |

<sup>1</sup>REML unbounded variance components ± standard error, Wald p-values

When raised under stress conditions, larval size at 75 dpf (i.e., after all larvae had hatched) was a significant predictor of juvenile size about 6 months later (LMM; F = 6.8, p = 0.01; Table S4a). The analogous correlation in the non-stressed controls was not significant (Table S4b). However, the stress-induced reduction in larval length at 75 dpf was no significant predictor of juvenile size in the wild (Table 4c).

Recapture rates per dam could not be predicted from average larval length (r_s_ = -0.31, n = 12, p = 0.32), yolk sac volume (r_s_ = 0.05, p = 0.88), stress-induced reduction in larval size (r_s_ = 0.32, p = 0.31), nor stress-induced difference in hatching time (r_s_ = -0.34, p = 0.28), and none of the analogous correlation for recapture rates per sire was significant (r_s_ < 0.30, n = 10, p always > 0.40). The significant maternal effects on juvenile body size that Bylemans et al. (2023) had found was confirmed in all LMM that included larval length or hatching date (Tables 1a; 2; S2, S3, S4), but was not significant in LMMs that included egg size, egg content, or yolk sac volume (Tables 1b, S1). There was never a significant sire effect on juvenile size in any LMM.

## Discussion

The high recapture rates that Bylemans et al. (2023) reported allowed, arguably for the first time, to statistically link a large-scale laboratory study on fish embryos (Wilkins et al. 2017) with the performance of their juvenile siblings in the wild. All parental fish were well represented among the recaptured juveniles. Bylemans et al. (2023) reported that the recapture rates could not be predicted from inbreeding coefficients. We found that the recapture rates were also not correlated to any traits that had been measured by Wilkins et al. (2017) in the laboratory. However, we found that juvenile growth in the wild could be well predicted by measures taken in the laboratory. Egg size, larval length, and yolk sac volumes were all significantly correlated with juvenile size (after taking possible parental effects into account). The significant dam effects in most of our statistical models confirm Bylemans et al. (2023)’s findings and suggest that other maternal environmental effects affected juvenile growth, too. Because Wilkins et al. (2017) had quantified egg carotenoid content, we could test for correlations between carotenoids and juvenile growth but found them not to be significant. It remains to be shown what other maternal environmental effects could play a role here. Females may differ, for example, in how they supply their eggs with innate immunity proteins and antibodies (Li and Leatherland 2012), and variance in maternal stress can influence glucocorticoid levels in eggs that then affect offspring development (Sopinka et al. 2017).

Wilkins et al. (2017) had experimentally challenged the embryos by exposing half of them to a bacterial pathogen that did not cause increased mortality but reduced growth and induced precocious hatching. This pathogen-related effect on hatching time was a good predictor of juvenile growth in the wild. Juvenile grew smaller if their siblings had shown a stronger reaction to the experimental stress in the laboratory than other sib groups. The analogous effects of pathogen-induced reduction in growth were not statistically significant, but the correlation between larval size and juvenile size seemed strongest in the pathogen-exposed group.

The timing of hatching has previously been demonstrated to be a good indicator of perceived stress in salmonids and other taxa. The nearby presence of an infected egg and even water-borne cues emitted from infected eggs can induce early hatching in brown trout (Pompini et al. 2013), other salmonids (Wedekind 2002), and even other lower vertebrates (Warkentin et al. 2001). Induced early hatching could also be observed in response to other types of stress such as simulated desiccation (Wedekind and Müller 2005), a simulated or actual attack by a predator (Warkentin 2005, Gomez et al. 2023), or exposure to chemical stressors (Lieke et al. 2021). The response of infected embryos can, however, differ in direction. Sometimes, exposure to pathogens induce early hatching (Warkentin et al. 2001, Wedekind 2002, Pompini et al. 2013), sometimes it delays hatching (Clark et al. 2014, Nusbaumer et al. 2021b). This difference in reaction is not understood yet but could be linked to the virulence of an infection.

Measuring family-specific fitness is notoriously difficult (Carlson and Seamons 2008) and often based on strong assumptions, especially in laboratory studies. We found that key variables that are typically used in laboratory studies on embryos, such as larval growth or stress-induced change in life history, were good predictors of how siblings of these embryos grew in the wild during their first spring and summer. Our findings support the implicit assumption of numerous studies, namely that the effects of environmental challenges, as measured under laboratory conditions, can serve as valuable predictors of family-specific performance in the wild and hence of the evolutionary potential of fish populations.

## Acknowledgements

We thank U. Gutmann and B. Bracher for catching and handling the adult fish, and for raising and stocking the F1 into the wild, L. Garaud, M. Hobil, C. Küng, E. Longange, L. Menin, D. Ortiz, E. Pereira-Alvarez, and V. Vocat-Mottier for help in the laboratory and/or for discussion, and T. Bösch, J. Kast, and the ecogenics team for genotyping the fish. The project was approved by the *Veterinärdienst* of the Bern canton (BE118/14) and by the fisheries inspectorate of the Bern canton. The Swiss National Science Foundation provided funding (31003A_159579 & 31003A_182265).

## Author contribution

JB, LMC, LW & CW designed the project. LMC, LW, DN, AU & CW did the experimental breeding, LMC & LW raised the embryos in the laboratory, and LMC, DN & AU sampled the juveniles from the wild. JB & LMC were responsible for the genotyping and parental assignments. JB & CW performed the analyses and wrote the manuscript. All authors revised and approved the final manuscript for publication.

## Supplementary Material

**Figure S1.**
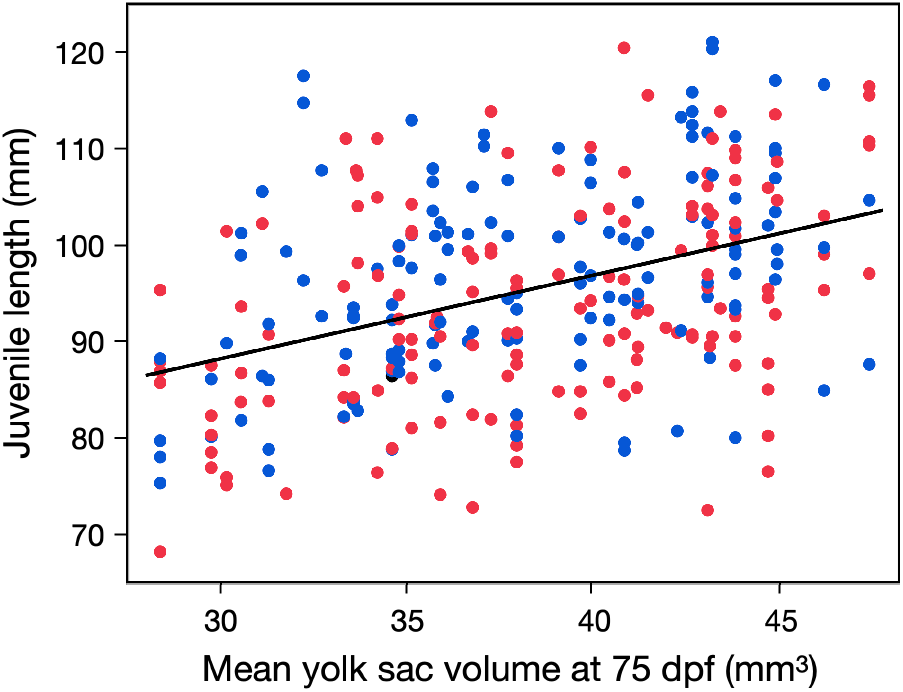
Juvenile length predicted by the mean yolk sac volume per full-sib family as determined about 6 months earlier at 75 dpf (days past fertilization), for male and female juveniles (blue and red symbols, respectively). The regression line is drawn to illustrate the direction of the correlation. See Table 1 for statistics that takes parental effects into account.

**Figure S2:**
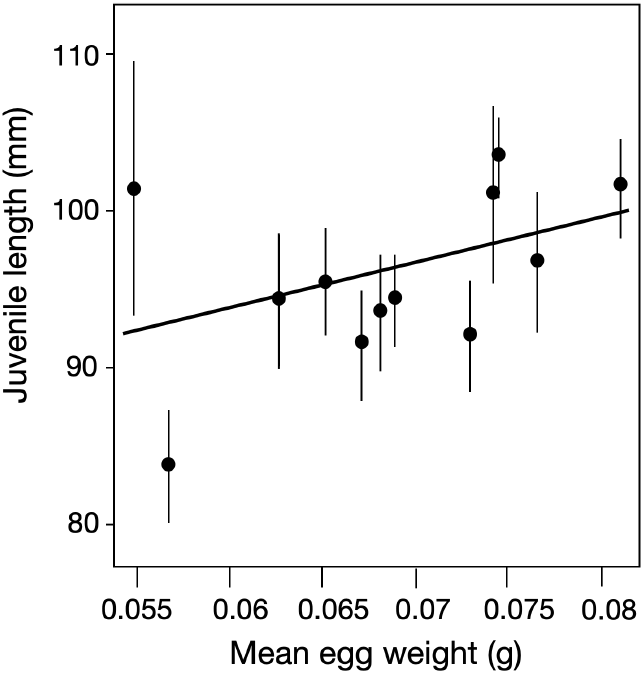
Mean (±95%CI) juvenile length (per maternal sib group) predicted by average egg weight. The regression line illustrates the direction of the correlation. See Table S1 for statistics.

**Table S1.** Linear mixed model on juvenile length as predicted by the egg characteristics. (a) Mean egg weight per full-sib family, and (b) the mean astaxanthin, capsanthin, and lutein content per egg. Zeaxanthin content was excluded from the model to avoid collinearity problems (the correlation between astaxanthin and zeaxanthin content was r = 0.56). Likewise, the effects of egg size and carotenoid content could not be tested in the same model because the correlation between egg weight and astaxanthin content was r = -0.67. Parental identities were included as random factors. Significant p-values are highlighted in bold.

| Effects | d.f. | F | Variance component | p |
| --- | --- | --- | --- | --- |
| *a) Egg weight* |  |  |  |  |
| Fixed effects: |  |  |  |  |
| Egg weight | 1 | 6.1 |  | **0.04** |
| Random effects<sup>1</sup>: |  |  |  |  |
| Dam |  |  | 15.7 ± 9.8 | 0.11 |
| Sire |  |  | 4.7 ± 3.7 | 0.20 |
| Residual |  |  | 81.8 ± 7.0 |  |
| *b) Carotenoid content* |  |  |  |  |
| Fixed effects: |  |  |  |  |
| Astaxanthin | 1 | 10.3 |  | 0.72 |
| Capsanthin | 1 | 7.8 |  | 0.19 |
| Lutein | 1 | 7.2 |  | 0.27 |
| Random effects<sup>1</sup>: |  |  |  |  |
| Dam |  |  | 31.3 ± 18.9 | 0.10 |
| Sire |  |  | 3.8 ± 3.4 | 0.26 |
| Residual |  |  | 81.4 ± 7.4 |  |

<sup>1</sup>REML unbounded variance components ± standard error, Wald p-values

**Table S2.**
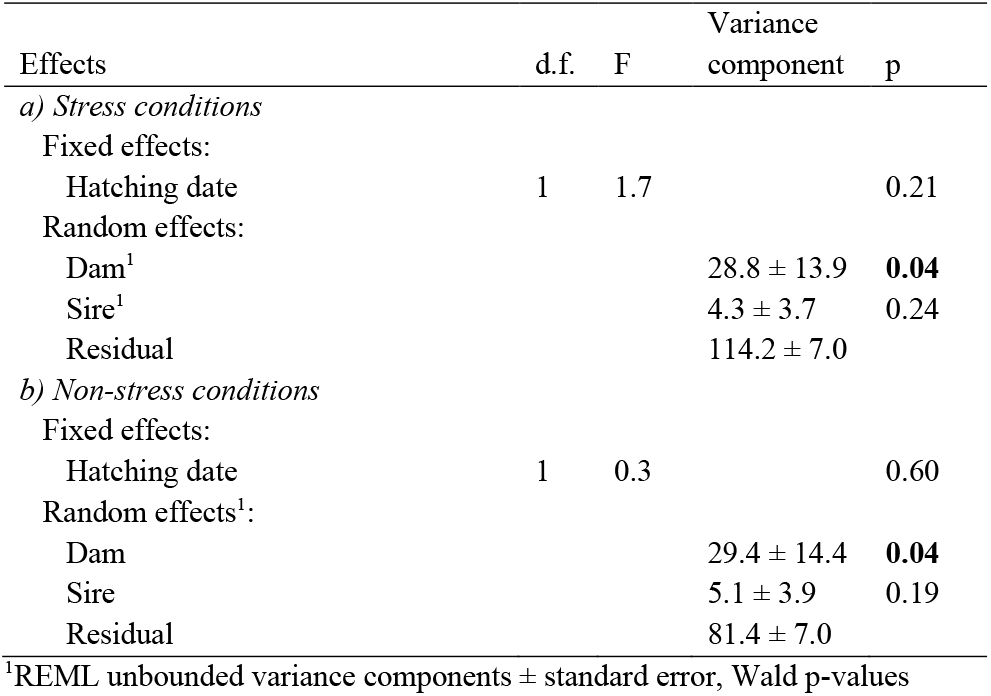
Linear mixed model on juvenile length as predicted by mean larval length at 75 dpf per family as measured under laboratory conditions about 6 months earlier under (a) stress conditions (exposure to *P. fluorescens*) or (b) non-stress conditions (controls). In (c), the stress-induced reduction in larval length (relative to larval length under control conditions) was used as possible predictor of juvenile length in the wild. Parental identities were included as random factors in all models. Significant p-values are highlighted in bold.

| Effects | d.f. | F | Variance component | p |
| --- | --- | --- | --- | --- |
| *a) Stress conditions* |  |  |  |  |
| Fixed effects: |  |  |  |  |
| Hatching date | 1 | 1.7 |  | 0.21 |
| Random effects: |  |  |  |  |
| Dam<sup>1</sup> |  |  | 28.8 ± 13.9 | **0.04** |
| Sire<sup>1</sup> |  |  | 4.3 ± 3.7 | 0.24 |
| Residual |  |  | 114.2 ± 7.0 |  |
| *b) Non-stress conditions* |  |  |  |  |
| Fixed effects: |  |  |  |  |
| Hatching date | 1 | 0.3 |  | 0.60 |
| Random effects<sup>1</sup>: |  |  |  |  |
| Dam |  |  | 29.4 ± 14.4 | **0.04** |
| Sire |  |  | 5.1 ± 3.9 | 0.19 |
| Residual |  |  | 81.4 ± 7.0 |  |

<sup>1</sup>REML unbounded variance components ± standard error, Wald p-values

**Table S3.**
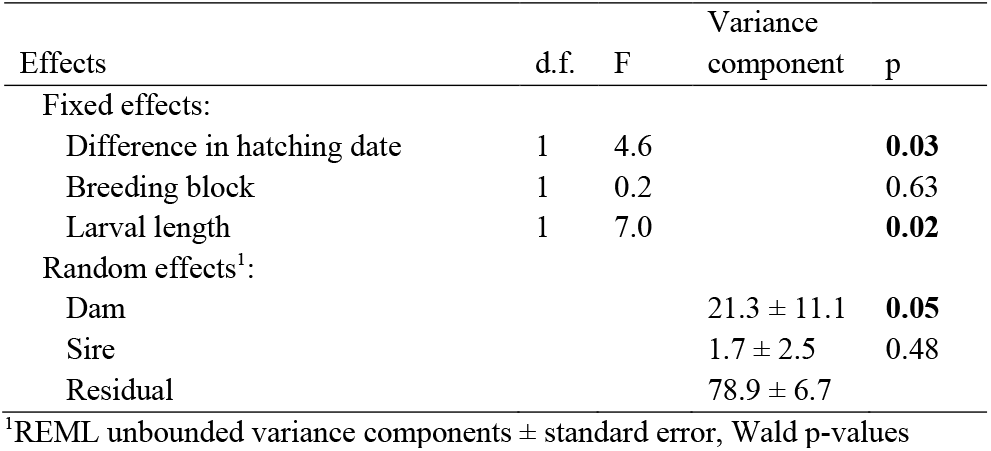
Linear mixed model on juvenile length as predicted by the stress-induced difference in hatching date under laboratory conditions, i.e., the difference between the mean hatching date under control conditions and the mean hatching date after exposure to *P. fluorescens*. The factors breeding blocks and mean larval length at 75 dpf (using means per family and treatment) were added as further fixed factors, while parental identities (nested in breeding block) were included as random factors. Significant p-values are highlighted in bold (after stepwise removal of non-significant interaction terms).

**Table S4.**
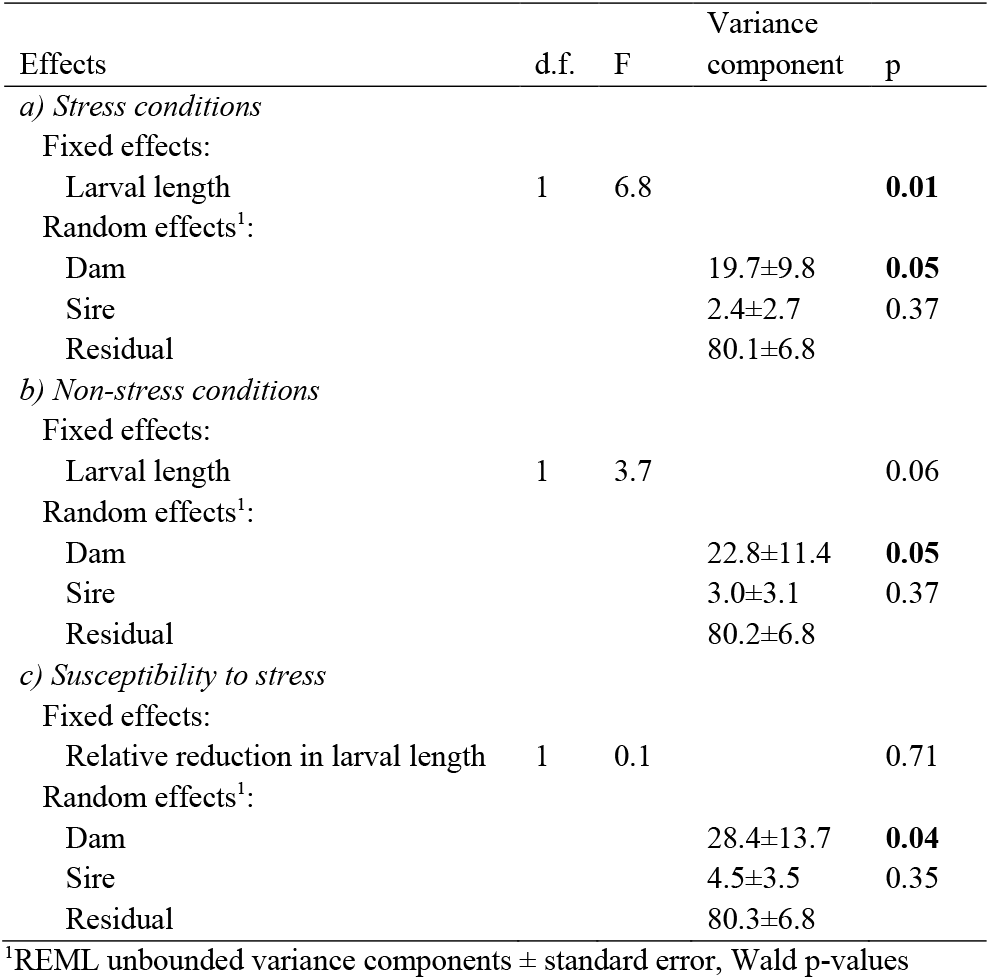
Linear mixed model on juvenile length as predicted mean larval length at 75 dpf per family as measured under laboratory conditions about 6 months earlier under (a) stress conditions (exposure to *P. fluorescens*) or (b) non-stress conditions (controls). In (c), the stress-induced reduction in larval length (relative to larval length under control conditions) was used as possible predictor of juvenile length in the wild. Parental identities were included as random factors in all models. Significant p-values are highlighted in bold.

| Effects | d.f. | F | Variance component | p |
| --- | --- | --- | --- | --- |
| *a) Stress conditions* |  |  |  |  |
| Fixed effects: |  |  |  |  |
| Larval length | 1 | 6.8 |  | **0.01** |
| Random effects<sup>1</sup>: |  |  |  |  |
| Dam |  |  | 19.7±9.8 | **0.05** |
| Sire |  |  | 2.4±2.7 | 0.37 |
| Residual |  |  | 80.1±6.8 |  |
| *b) Non-stress conditions* |  |  |  |  |
| Fixed effects: |  |  |  |  |
| Larval length | 1 | 3.7 |  | 0.06 |
| Random effects<sup>1</sup>: |  |  |  |  |
| Dam |  |  | 22.8±11.4 | **0.05** |
| Sire |  |  | 3.0±3.1 | 0.37 |
| Residual |  |  | 80.2±6.8 |  |
| *c) Susceptibility to stress* |  |  |  |  |
| Fixed effects: |  |  |  |  |
| Relative reduction in larval length | 1 | 0.1 |  | 0.71 |
| Random effects<sup>1</sup>: |  |  |  |  |
| Dam |  |  | 28.4±13.7 | **0.04** |
| Sire |  |  | 4.5±3.5 | 0.35 |
| Residual |  |  | 80.3±6.8 |  |

<sup>1</sup>REML unbounded variance components ± standard error, Wald p-values

